# Use of RNA and DNA to Identify Mechanisms of Microbial Community Homogenization

**DOI:** 10.1101/496679

**Authors:** Kyle M. Meyer, Ian A. B. Petersen, Elie Tobi, Lisa Korte, Brendan J. M. Bohannan

**Author notes:** Address correspondence to Kyle M. Meyer,.

## Abstract

Biotic homogenization is a commonly observed response following conversion of native ecosystems to agriculture, but our mechanistic understanding of this process is limited for microbial communities. In the case of rapid environmental changes, inference of homogenization mechanisms may be confounded by the fact that only a minority of taxa is active at any given point. RNA- and DNA-based community inference may help to distinguish the active fraction of a community from inactive taxa. Using these two community inference methods, we asked how soil prokaryotic communities respond to land use change following transition from rainforest to agriculture in the Congo Basin. Our results indicate that the magnitude of community homogenization is larger in the RNA-inferred community than the DNA-inferred perspective. We show that as the soil environment changes, the RNA-inferred community structure tracks environmental variation and loses spatial structure. The DNA-inferred community loses its association with environmental variability. Homogenization of the DNA-inferred community appears to instead be driven by the range expansion of a minority of taxa shared between the forest and conversion sites, which is also seen in the RNA-inferred community. Our results suggest that complementing DNA-based surveys with RNA can provide unique perspectives on community responses to environmental change.

**IMPORTANCE:** Two primary mechanisms by which community homogenization occurs are: 1) the loss of environmental heterogeneity driving community convergence, and 2) increased rates of biotic mixing, driven by exotic invasions or range expansions. Better identifying these mechanisms could help inform future mitigation strategies. Only a minority of soil taxa tends to be active at any time, which makes identifying these mechanisms difficult. To circumvent this problem, we measured prokaryotic community structure in two ways: RNA-based inference (which should enrich for active taxa), and DNA-based inference (which includes active and inactive taxa) along a gradient of land use change. Our results suggest that changes to soil heterogeneity impact the RNA-inferred community, while range expansions contribute to the homogenization of both DNA- and RNA-inferred communities. Thus, RNA-based community inference may be a more sensitive indicator of environmentally driven homogenization, and researchers interested in microbial responses to rapid environmental change should consider this method.

## INTRODUCTION

One of the most rampant forms of environmental change today is land use change following the conversion of tropical rainforests to agriculture (1–4). Both above- and below-ground communities have been shown to experience species loss and community change at unprecedented rates following land use change (5–7), and this is of concern because tropical rainforests are some of the most diverse and productive ecosystems on the planet. Predicting community responses to tropical land use change is a priority if we are to better understand how human activities will impact species loss and global-scale biogeochemical cycling (8, 9), but in order to gain such a level of predictability we must better understand the mechanisms underlying community change.

Biotic homogenization, *i.e*. the increase in community similarity through time or space, is a major consequence of land use change (10, 11). This process can be driven by two primary mechanisms: 1) the loss of environmental heterogeneity, which drives subsequent community convergence (12, 13), and 2) increased rates of biotic mixing, which can be driven by the breakdown of dispersal barriers, invasion of exotic taxa, or the range expansion of existing taxa (11, 14). These mechanisms have both been implicated in the homogenization of microbial communities following land use change (6, 7, 15–18), but it remains unclear to what degree these mechanisms contribute to homogenization.

Understanding mechanisms of biotic homogenization may be complicated by the fact that only a minority of soil taxa tends to be active at any given point in time (19, 20). One proposed method to distinguish active community members is to survey the community using 16S rRNA (as opposed to the 16S rRNA gene) (21–24). This methodology could provide new insights into microbial community homogenization. For example, targeting active taxa could help us hone in on the portion of the community that is interacting with the environment and thus who is likely to respond immediately to environmental changes. Secondly, if land use change is driving increased rates of biotic mixing, studying the active fraction could help us distinguish who is actually growing and becoming established from those who are simply arriving. This distinction may be especially important when considering that much of what we currently know about microbial homogenization has been derived from DNA-based diversity studies (e.g. (6, 25, 27)) that do not distinguish active from inactive taxa. Some controversy, however, surrounds the use of rRNA to infer microbial activity levels. For example, rRNA concentration and growth rate and/or activity are not consistently correlated across taxa, and certain taxa can still contain ribosomes while dormant (see (28)). The use of 16S rRNA: 16S rRNA gene ratios of taxa has also been shown to not correlate well with activity levels inferred by other means (29, 30), and can be biased by extracellular environmental DNA (31), taxon-specific dormancy strategies and sampling extent (32). While the use of rRNA:rDNA ratios may be problematic, several studies have shown that communities inferred using rRNA more closely correlate with environmental variability (33), and respond more strongly to seasonal variation (34) and nutrient pulses (35) than communities inferred using rDNA, which is consistent with the idea that the rRNA content of a community is at least enriched with active members. Thus, RNA-based community inference may provide unique foundational insights into the mechanisms underlying community change, but to date few have sought to make this comparison.

Despite growing efforts to characterize microbial responses to land use change, a number of fundamental gaps must be filled to bring our understanding to a more generalizable level. For example, although there have been several studies comparing established agricultural sites to pristine ecosystems, few have sought to include sites that represent the intermediary stages of conversion (*e.g*. recently slash-and-burned areas). By including more sites along the conversion continuum, we can increase the resolution by which we understand this process. This could help to diagnose when the largest losses of biodiversity occur, and pinpoint management practices that could be targeted for improvement. Another important gap to fill lies in the geographic representation of sampling efforts. By expanding sampling efforts geographically, we can start to distinguish common patterns from site-specific patterns. This is especially important when considering that much of what we know about microbial responses to tropical land use change comes from studies in the Amazon Basin (6, 7, 15, 25, 36–42), and to a lesser degree, the forests of Indonesia (16, 17, 26, 43) with far fewer studies in the forests of Central and West Africa (27, 44, 45), and to our knowledge, none in the Congo Basin. Thus, by focusing our efforts to study the conversion process with more resolution and a wider geographic representation, we can work towards a more generalizable understanding of microbial responses to tropical land use change.

Here we examine soil bacterial community change along a land use change gradient in the Congo Basin, the world’s second largest rainforest (46). Our work expands on past studies by performing paired RNA/DNA co-extraction from each sample in order to ask whether the putatively active fraction of the community elicits a different response to land use change than the total community. Our gradient includes a site that had very recently been cut and burned, which allows us to use RNA/DNA in a system that is experiencing rapid and intense change. We test the following hypotheses: 1) that converted (burned and plantation) sites will exhibit decreased rates of spatial turnover of both the RNA- and DNA-inferred prokaryotic communities, 2) that changes to the soil chemical environment will play a stronger role in shaping the RNA-inferred community than the DNA-inferred community, and 3) that biotic invasions or range expansions contribute to community homogenization.

## MATERIALS & METHODS

### Sampling site

Central Africa contains up to 1.8 million km^2^ of contiguous tropical moist forest, making it the second largest block of tropical moist forest in the world, after the Amazon Basin (46). Central African rainforest is renowned for its exceptionally high levels of biodiversity and endemism (47–49) and it is rapidly being deforested (50). The nation of Gabon contains more than 10% of the contiguous tropical moist forest in Africa (46, 47), and the majority of these forested areas are either currently leased as long-term logging concessions or are at risk from agricultural conversion (47, 51, 52).

Our study was performed in southwestern Gabon near the Gamba Complex of Protected Areas (47). Soils in this area are classified as Dystic Fluvisol (53). Agricultural conversion in this region follows slash-and-burn practices that are typical of most tropical regions whereby forests are selectively logged and the remaining vegetation is burned. The following season, plantation crops (typically manioc or banana) are planted and harvested for 1-3 years. Following the last harvest, plantations are abandoned and secondary forest develops. We selected sites representative of this cycle including a recently burned site, an active manioc and banana plantation (roughly 1.5 years old), and an adjacent intact forest, which allows us to break down the conversion process into two steps, providing more resolution. Sites are found at the following coordinates: burned site (2° 44′ 48″ S, 10° 8′ 54″ E), plantation (2° 44′ 58″ S, 10° 8′ 51″ E), and adjacent forest (2° 44′ 46″ S, 10° 8′ 52″ E).

### Sampling Design and Sample Collection

This study was designed specifically to understand differences between RNA- and DNA-inferred communities within these sites, not to identify general effects of land use change on Congo Basin ecosystems, which would be better-tested using replication at the land type level (54). Limited access to sites and logistical challenges with sampling in this area required that we extensively survey one site within each of three land types, rather than performing higher levels of replication on fewer land types. This design is appropriate for asking how these sites differ from one another, or how RNA- and DNA-inferred community composition or diversity patterns differ from one another (55–57). Regarding inferences about general microbial responses to land use change in the Congo Basin, this study would be considered a case study (54), whereby our results may be suggestive of broader patterns, but such patterns should be corroborated using a design with land type replication.

Soil samples were taken at the end of the Gabonese dry season (September 24-27, 2013). We established plots within each of the aforementioned sites. Each plot consisted of a nested sampling scheme (6) where a 100 m x 100 m quadrat was established, with 10 m x 10 m, 1 m x 1 m, 0.1 m x 0.1 m quadrats nested within each, giving high coverage of a range of spatial scales (Fig. 1). Soil cores were taken to a depth of 15 cm (after removal of leaf litter) from the corners of each quadrat (N=13 samples per site). For each point, 3 cores were taken, homogenized, and then subsampled. From the homogenized mixture, 3 ml (approximately 1 g) of soil was added to 9 ml Lifeguard solution (Mobio, California, USA) in the field, then transported cold and stored at −80° C in order to stabilize nucleotides for later extraction. Our spatially explicit design allows for the estimation of spatial turnover (beta diversity)(58).

**Figure 1:**
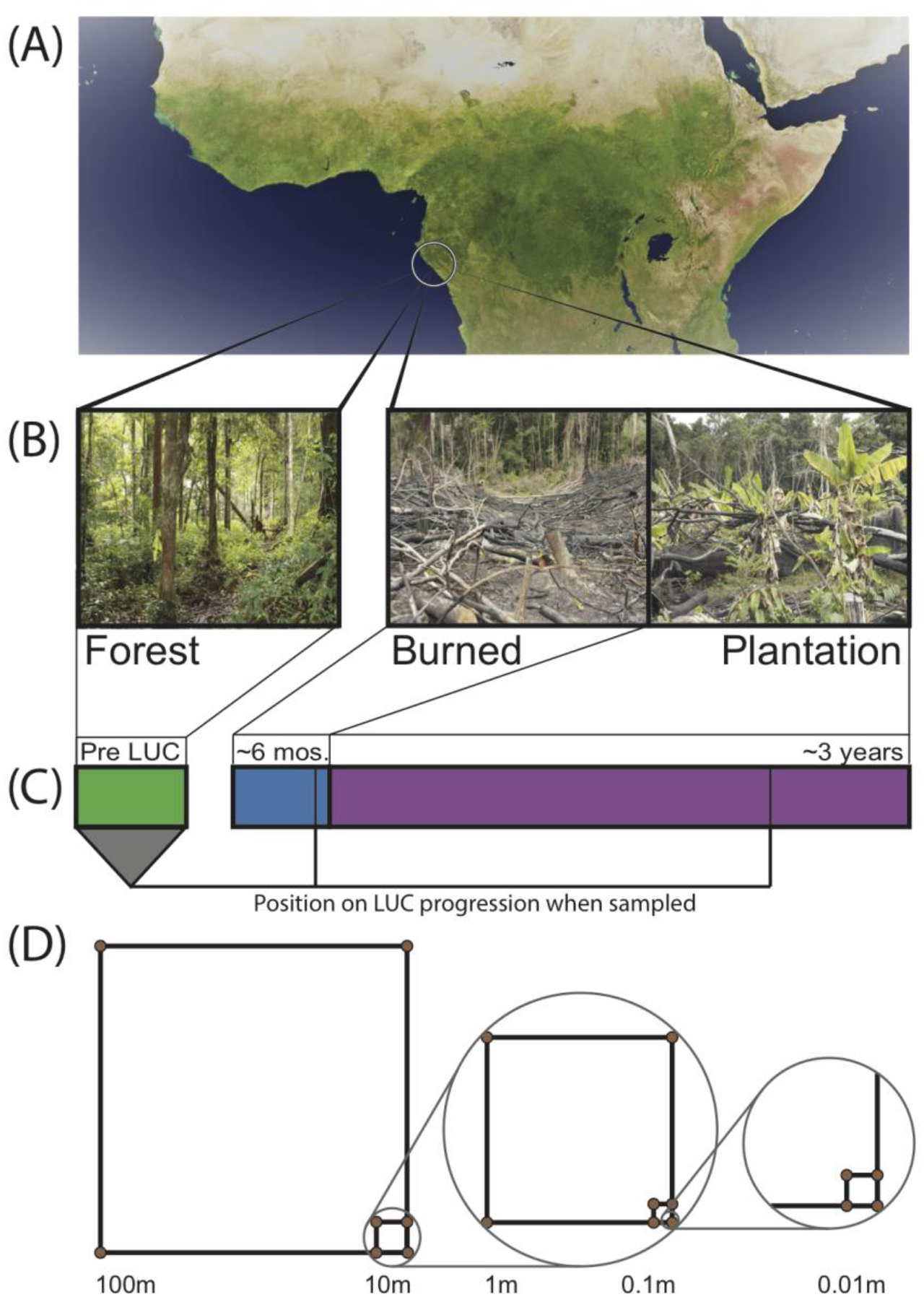
Sampling design across Gabonese chronosequence of land use change. A) Satellite image of the Congo Basin with location of sampling sites circled. B) Images of field sites from which samples were taken. C) Timeline of land use change. Bar width is proportional to the amount of time a site typically spends in each stage. Lines indicate when samples were collected. D) Spatially explicit nested sampling scheme used in each land type. Samples were taken at the corners of each square.

### Extraction, PCR, and Sequencing

Soil RNA and DNA were co-extracted from Lifeguard-preserved soil samples using MoBio’s Powersoil RNA Isolation kit with the DNA Elution Accessory Kit (MoBio, California, USA) following manufacturer’s instructions. Extractions were quantified using Qubit (Life Technologies, USA). RNA was reverse transcribed to cDNA using Superscript III first-strand reverse transcriptase and random hexamer primers (Life Technologies, USA).

The V3 and V4 region of the 16S rRNA gene of the DNA and cDNA were PCR amplified using the primers 319F and 806R (primarily targeting Bacteria, with limited coverage of Archaea). Sequencing libraries were prepped using a two-step PCR with dual-indexing approach (59, 60). In short, the first round of amplification consisted of 22 cycles with Phusion HiFi polymerase. Round 1 products were cleaned using Agencourt AMPure XP (Beckman Coulter, California, USA) then amplified for an additional 6 cycles using Phusion HiFi to add the sequences required for cluster formation on the Illumina flowcell. The final library was sent to the Dana-Farber Cancer Institute Molecular Biology Core Facilities for 300 paired-end (PE) sequencing on the Illumina MiSeq platform.

### Soil chemical analysis

Soil chemical parameters were measured in each soil core (by A & L Western Agricultural Lab, Modesto, CA, USA), including percent organic matter (loss on ignition (61)), extractable phosphorus (Weak Bray (62) and sodium bicarbonate (63)), extractable cations (K, Mg, Ca, Na, by ammonium acetate extraction (64)), nitrate-N, sulfate-S (65), pH, buffer pH, cation exchange capacity (CEC, (66)), and percent cation saturation.

Pearson’s correlation tests were performed on all pairs of chemical parameters to test for autocorrelation and reduce the number of chemical variables used in our models. Pairs of variables that were highly correlated (R^2^ > 0.6, *P* < 0.05) were reduced to a single variable. The final suite of chemical analyses used after paring down correlated variables included percent organic matter, extractable phosphorus (Weak Bray), pH, extractable K, CEC, nitrate-N, and S.

### Bioinformatics and statistical analysis

Paired end reads were joined then demultiplexed in QIIME (67) before quality filtering. Primers were removed using a custom script. UPARSE was used to quality filter and truncate sequences (416bp, EE 0.5) (68). Sequences were retained only if they had an identical duplicate in the database. Operational taxonomic units (OTUs) were clustered *de novo* at 97% similarity using USEARCH (69). OTUs were checked for chimeras using the gold database in USEARCH. We used a custom script to format the UCLUST output for input into QIIME. To assign taxonomy, we used the repset from UPARSE in QIIME using greengenes version 13_5 (RDP classifier algorithm). Finally, we averaged 100 rarefactions at a depth of 3790 counts per sample for each community inference (RNA or DNA) and each land type (forest, burned, or plantation) to achieve approximately equal sampling depth across comparisons, which excluded three samples in the DNA-inferred communities (two in the forest and one in the plantation).

Statistical analyses were performed in the R platform (70). Canberra pairwise community distances were calculated using the vegdist function in the package ‘vegan’ (71). Canberra was chosen because of its incorporation of abundance data, sensitivity to rare community members (72), and ability to detect ecological patterns even in instances of relatively low sampling extent (73). Rates of community spatial turnover were estimated by regressing pairwise community similarity (1- Canberra distance) against pairwise geographic distance between samples (74). We used a similar regression approach between community similarity and environmental similarity to estimate the relationship between community turnover and environmental turnover. Pairwise soil environmental similarity was calculated using 1- Gower dissimilarity (75, 76) using the daisy function in the package ‘cluster’ in R (77). Gower dissimilarity was chosen because it can incorporate and compare different classes or scales of data (78). Mantel tests were used to test for significant associations between geographic, community, and environmental distance, and partial Mantel tests were used to estimate the relative contribution of environmental distance and geographic distance on variation in community dissimilarity in the ‘vegan’ package in R. Differences in average pairwise similarity across land types were assessed using a one-way ANOVA after verifying normal distribution of data. Post-hoc comparisons of group means were made using Tukey’s HSD. Distance-decay slopes were compared using the function diffslope (package ‘simba’) (79). This function employs a randomization approach across samples from each dataset and compares the difference in slope to the original configuration of samples. The p-values computed are the ratio between the number of cases where the differences in slope exceed the difference in slope of the initial configuration and the number of permutations (1000). We used the DESeq2 function (80) in R to identify differentially abundant taxa in one land type versus another. Low abundance samples were excluded prior to performing DESeq2 analysis. This function uses a generalized linear model (family negative binomial) to estimate dispersion and log2-fold change in relative abundance of individual taxa. Taxa were deemed differentially abundant if they had a positive log_2_-fold change and *P_adj_* < 0.05. Figures were either created using base R or the ‘ggplot2’ package (81).

We developed several community analysis approaches to investigate whether biotic invasion or range expansion contribute to biotic homogenization. Taxa found in a conversion land type (i.e. the burned or plantation site), but not the forest, were considered “newcomers”. We removed these taxa from the community matrix, equalized sampling extent (using rarefaction), and then re-ran analyses of pairwise community similarity levels and distance-decay (described above). The expectation was that if they contribute to homogenization (increased community similarity), then their removal should decrease pairwise community similarity levels. We took an analogous approach to ask if range expansion of forest-associated taxa (referred to as “bloomer” taxa) contributes to biotic homogenization. We identified taxa that were differentially abundant in converted sites relative to the forest site (described above), then removed them from the community matrix of the converted site and re-assessed community similarity levels and distance-decay. The expectation, as above, was that if these taxa contribute to homogenization, then their removal should render the communities less similar.

### Data availability

DNA and cDNA sequence FASTA files, OTU tables, soil environmental data, as well as the R script for analysis will be available for download from 10.6084/m9.figshare.5930434.

## RESULTS

### Soil bacterial community structure differs by land use and community inference method

We first asked whether bacterial community structure differed by land use or by community inference method (*i.e*. RNA- or DNA-inference) by performing a PERMANOVA on OTU-level community Canberra distance, with land type and inference method as the dependent variables. Both variables were significant (land type F_2,73_ = 3.67, R^2^ = 0.089, *p* < 0.001, community inference method F_1,73_ = 4.70, R^2^ = 0.057, *p* < 0.001), indicating that bacterial communities differ in membership across sites, and that RNA- and DNA-inferred communities differ in membership. These findings were also consistent at higher taxonomic levels (Supp. Figs 1, 2, & 3). The most pronounced differences at the phylum level were lower relative abundances of Acidobacteria in the burned site compared to the forest and plantation sites (burned site (DNA): 6.86 +/− 0.78%, forest site (DNA): 11.07 +/− 1.73%, plantation site (DNA): 11.30 +/− 1.32%), and higher relative abundances of Actinobacteria in the burned relative to forest and plantation sites (burned site (DNA): 10.86 +/− 1.16 %, forest site (DNA): 7.69 +/− 1.40%, plantation site (DNA): 8.73 +/− 1.33%), and this trend was consistent whether communities were inferred via DNA or RNA (Supplemental Fig. 1). OTU-level richness also differed by land type (F_2,70_ = 8.26, *p* < 0.001), but not community inference method (p=0.80), with the burned site being significantly lower in richness than the forest or plantation sites (Tukey’s HSD *p* < 0.01, for both comparisons, Supp. Fig. 4).

### Evidence of biotic homogenization following land use change

We asked whether soil prokaryotic communities in the sites undergoing agricultural conversion were on average more similar to each other, relative to the communities found in the forest. The RNA-inferred community showed a strong trend towards homogenization across sites (F_2,219_ = 23.33, *p* < 0.001, Fig. 2A), with average pairwise similarity progressively increasing over the chronosequence (1- Canberra dissimilarity: forest mean: 0.289 +/− 0.008, burned mean: 0.327 +/− 0.004, plantation mean: 0.340, +/− 0.004). The DNA-inferred community also differed in pairwise similarity across sites (F_2,184_ = 4.54, *p* = 0.012, Fig. 2B), but this trend was less pronounced, and similarity levels were only significantly higher in the burned site (1- Canberra dissimilarity: forest mean: 0.268 +/− 0.011, burned mean: 0.301 +/− 0.006, plantation mean: 0.288 +/− 0.006).

**Fig. 2:**
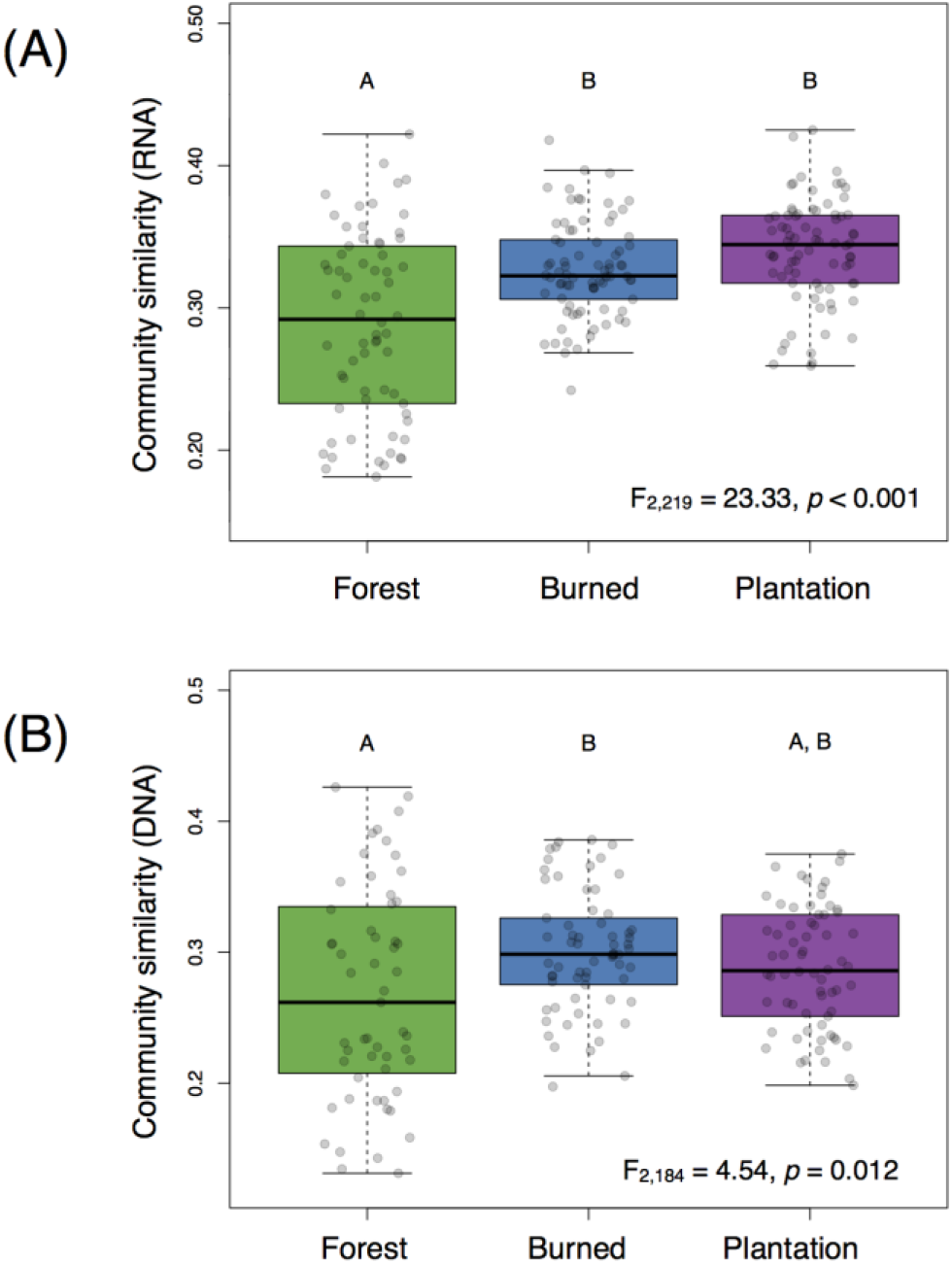
Average pairwise similarity (1- Canberra distance) of A) the RNA-inferred community, and B) the DNA-inferred community, across the forest, burned, and plantation sites. F and *p* statistics based on one-way ANOVA. Different letters correspond to significantly different group means as determined by Tukey’s HSD *p* < 0.05.

While levels of average pairwise community similarity tended to increase across the chronosequence, the spatial signal of community similarity (i.e. spatial turnover) tended to either weaken or disappear. Both the RNA-inferred and DNA-inferred communities showed distance-decay relationships in the forest (Mantel r_RNA_ = 0.846, *p* = 0.003, slope = −0.027; Mantel r_DNA_ = 0.697, *p* = 0.02, slope = −0.028, Fig. 3A,B) where communities in close proximity tended to exhibit higher levels of similarity than communities farther apart. The RNA-inferred community showed no significant distance-decay relationship in either the burned (Mantel r = 0.247, *p* = 0.127) or the plantation (Mantel r = 0.431, *p* = 0.063) sites. The DNA-inferred community showed a weak distance-decay relationship in the burned site with a three-fold decrease in slope from the forest (Mantel r = 0.474, *p* = 0.048, slope = −0.009), and no significant distance-decay relationship in the plantation (Mantel r = 0.232, *p* = 0.163). Thus, both windows into the community indicated shifts towards spatial homogenization, but this trend was more pronounced in the RNA-inferred fraction of the community.

**Fig. 3:**
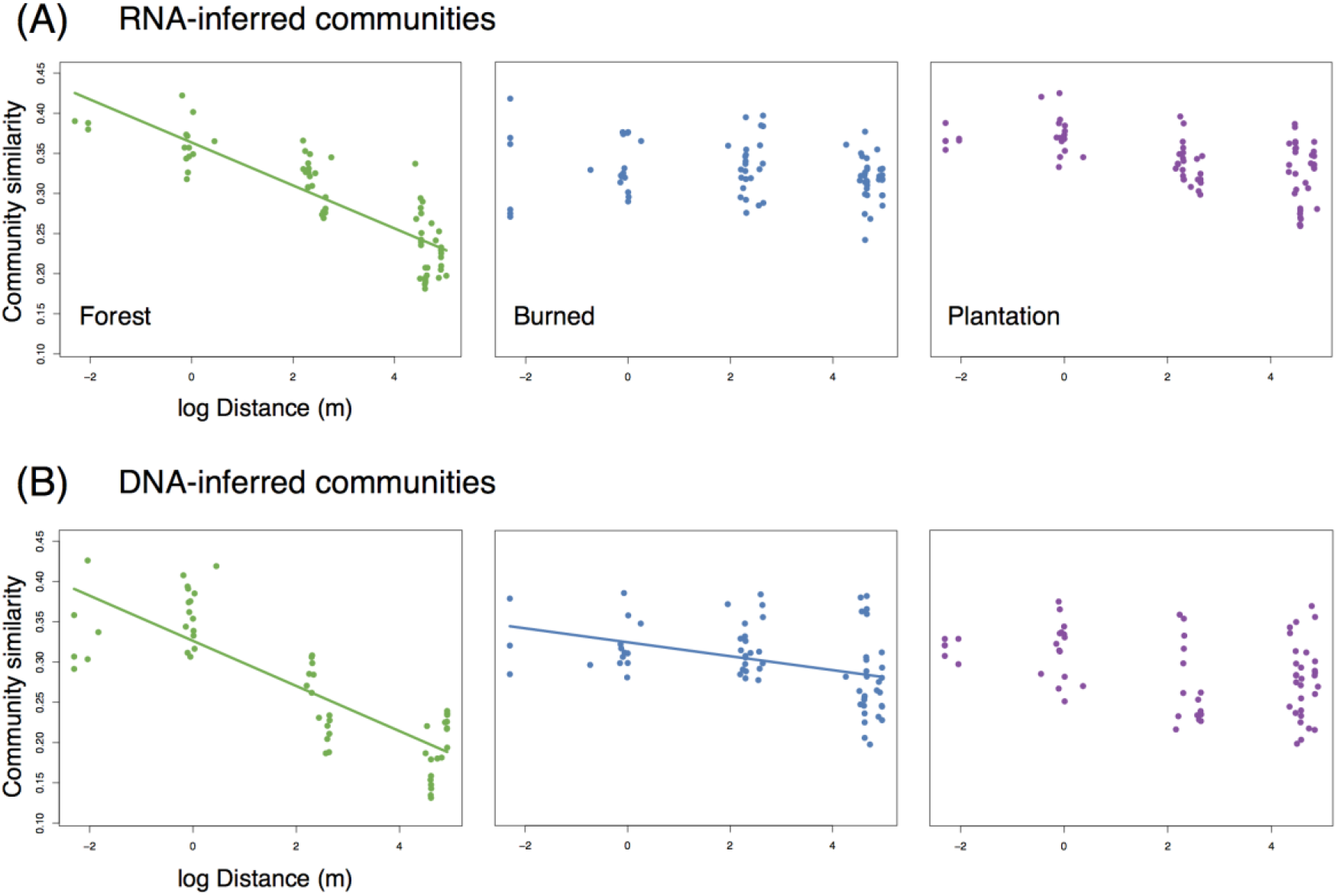
Change (or loss) of distance-decay of community similarity for A) RNA-inferred communities, and B) DNA-inferred communities. Trend lines were drawn only for significant (Mantel *p* < 0.05) associations.

### Soil environment gains variation, but loses spatial structure following conversion

Soil chemical profiles exhibited a number of changes across land types including increases in pH and phosphorus and decreases in percent organic matter throughout the chronosequence, and elevated cation exchange capacity and levels of nitrate-N, sulfur, and potassium in the burned site (Supplementary Table 1). When we consider the differentiation of soil chemical profiles within land types, we see that levels of average environmental pairwise similarity (1-Gower distance) decrease from the forest to the burned and plantation sites (F_2,231_ = 4.22, *p* = 0.016, Supplementary Fig. 5), indicating that soils within a land type are more dissimilar from one another. Similar to the spatial structure of the communities, the spatial structure of environmental variation also changes across the chronosequence. Forest soils show a significant environmental distance-decay relationship (Mantel r = 0.729, *p* = 0.01, slope = −0.052), where samples closer in proximity tend to be more similar in environmental conditions. This relationship was not significant in the burned site (Mantel r = 0.338, *p* = 0.068), and was comparatively weaker in the plantation relative to the forest (Mantel r = 0.465, *p* = 0.01) and showed a shallower distance-decay slope (slope = −0.027, difference in slope = −0.025, *p* = 0.001). Thus burning and planting seem to introduce environmental heterogeneity, but this heterogeneity tends to show little to no spatial structure.

### Environmental heterogeneity continues to influence RNA-inferred (and not DNA-inferred) community turnover, despite loss of spatial structure

We asked whether the loss of spatial structure of the soil chemical environment could be contributing to the loss of spatial turnover in the microbial community. To do so, we regressed pairwise community similarity (1-Canberra distance) against pairwise environmental similarity (1-Gower distance) for both the RNA- and DNA-inferred communities. In the forest site, both RNA- and DNA-inferred community similarity levels were positively correlated with environmental similarity (Fig. 4A, B), even after accounting for differences due to geographic distance (Table 1), suggesting samples with similar environmental (chemical) conditions tended to harbor similar communities. When we look at the burned and plantation sites, however, this relationship persists for the RNA-inferred community, but disappears for the DNA-inferred community (Table 1), suggesting that the spatial homogenization of the DNA-inferred community may be driven by other mechanisms besides soil chemical homogenization. Thus as environmental heterogeneity loses its spatial structure, the RNA-inferred community similarity levels continue to vary with this heterogeneity and lose spatial structure, while the DNA-inferred community becomes decoupled from levels of environmental variation.

**Fig. 4:**
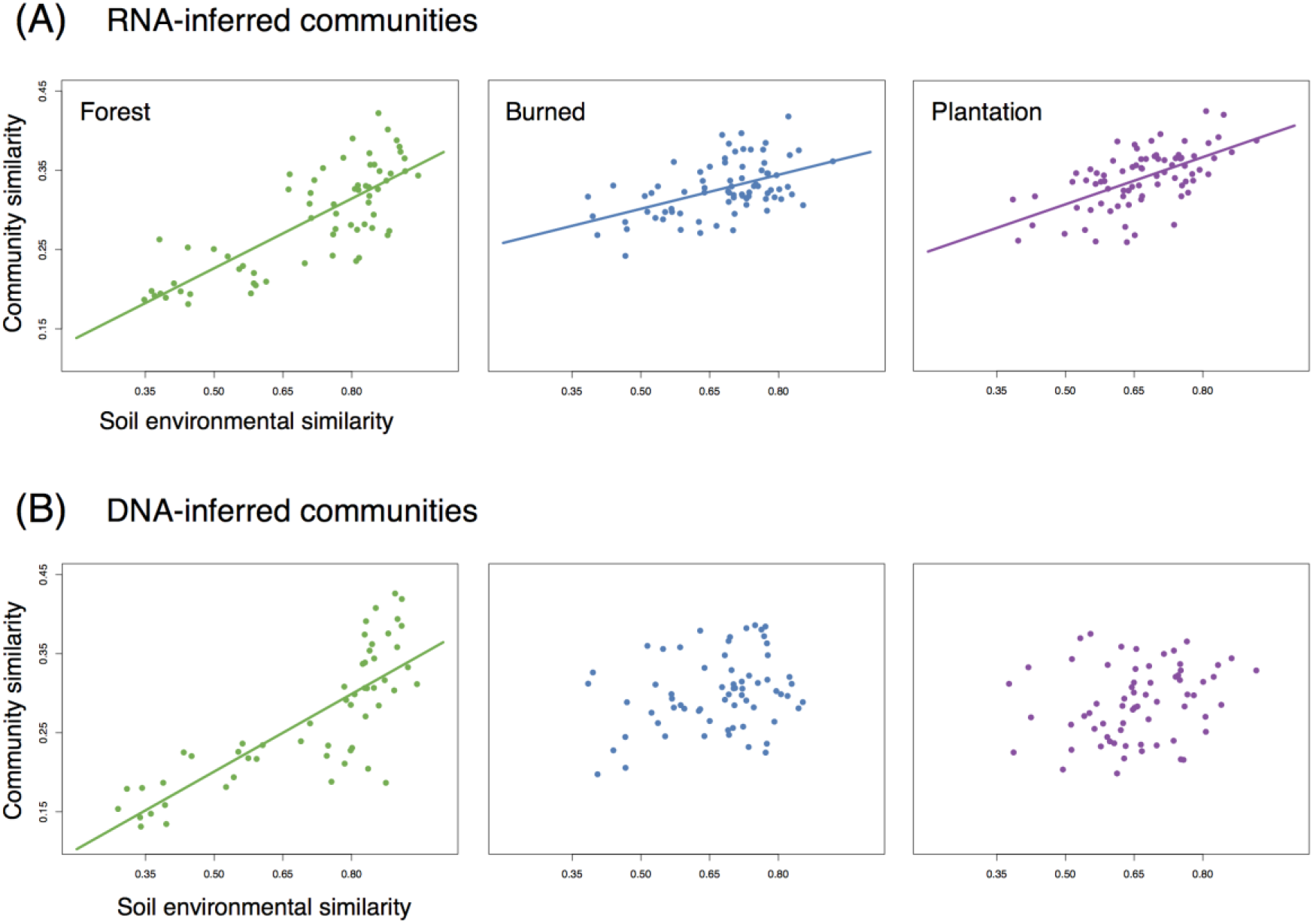
The relationship between community similarity and environmental similarity (1- Gower dissimilarity) for A) RNA-inferred communities, and B) DNA-inferred communities. Trend lines were only drawn for significant (Mantel *p* < 0.05) associations.

**Table 1:** The influence of environmental similarity and geographic distance on RNA-inferred and DNA-inferred prokaryotic communities. Partial mantel test summary statistics showing 1) the effect of environmental similarity after removing the effect of geographic distance (Env. Simil.), and 2) the effect of geographic distance after removing the effect of environmental similarity (Geog. Dist.). *P* values estimated from 1000 permutations.

|  | Env. Simil. |  | Geog. Dist. |  |
| --- | --- | --- | --- | --- |
|  | *r* | *P* | *r* | *P* |
| **Forest RNA** | **0.56** | **0.022** | 0.409 | 0.088 |
| **Burned RNA** | **0.491** | **0.008** | 0.05 | 0.392 |
| **Plantation RNA** | **0.57** | **0.001** | 0.082 | 0.284 |
| **Forest DNA** | **0.536** | **0.026** | 0.168 | 0.208 |
| Burned DNA | 0.237 | 0.174 | 0.338 | 0.07 |
| Plantation DNA | 0.17 | 0.194 | 0.124 | 0.267 |

### Biotic invasions do not contribute to homogenization

We next tested the hypothesis that the introduction of “newcomer” taxa (i.e. those that were not previously present) was driving community homogenization. 390 of the 1545 DNA-inferred community members in the burned site (25.2% of OTUs, representing on average 1.8 +/− 1.1% of the community) were not detected in the forest site. 570 of the 1804 DNA-inferred community members in the plantation site (31.7% of OTUs, representing on average 1.8 +/− 0.8% of the community) were not detected in the forest site. These taxa were not particularly geographically widespread (average occurrence frequency_Burn newcomers_ = 0.23 +/− 0.010, frequency_Plantation newcomers_ = 0.25 +/− 0.010). Moreover, only 53.6% of the newcomer taxa in the burned site DNA-inferred community were detected in the RNA fraction of that site, and 63.5% of the newcomer taxa in the plantation site DNA-inferred community were detected in the RNA fraction of that site, suggesting that not all newcomers may become established. We tested whether the newcomer taxa were driving higher estimates of community similarity by removing them from the community matrix, equalizing sampling extent across samples (using rarefaction), then re-calculating community similarity (1-Canberra distance). Our expectation was that the removal of newcomers from the community matrix would render communities more dissimilar (i.e. less homogenized). This was not the case. Removal of the newcomer taxa from the burned site community matrices actually increased community similarity of the DNA-inferred community (0.322 +/− 0.006 vs 0.301 +/− 0.006). Removal of newcomers also did not render a significant spatial signal for the DNA-inferred communities (Mantel_no newcomers_ r = 0.407, *p* = 0.074). This was also the case for the plantation where the removal of the newcomer taxa increased community similarity for the DNA-inferred communities (0.322 +/− 0.006 vs 0.288 +/− 0.006), and left no spatial signal (Mantel _no newcomers_ r = 0.220, *p* = 0.159). Thus we have no evidence to suggest that the homogenization of the DNA-inferred community is driven by the arrival of newcomer taxa.

We found similar results when we performed these same analyses on the RNA-inferred communities. Newcomer taxa comprised a similarly small proportion of the communities in the burned (2.2 +/− 1.8%) and plantation sites (2.1 +/− 0.79%). These taxa were also not particularly widespread, with average occurrence frequencies in the burned site of 0.26 +/− 0.16 and 0.25 +/− 0.01 in the plantation site. Similar to the DNA-inferred community, the removal of newcomer taxa from the RNA-inferred community rendered higher levels of average pairwise similarity in the burned site community (0.347 +/− 0.004) and the plantation site community (0.372 +/− 0.004), suggesting that their abundances are not likely increasing levels of community similarity. Lastly, the removal of newcomers from the RNA-inferred community did not render significant relationships with geographic distance (Burned site: Mantel _no newcomers_ r = 0.231, *p* = 0.151, Plantation site: Mantel _no newcomers_ r = 0.452, *p* = 0.07), indicating that they likely do not play a role in community spatial homogenization. Hence we have no evidence to support the hypothesis that increased levels of biotic homogenization are being driven by the arrival of newcomer taxa, and in fact, it appears that the newcomers may actually contribute variation to the communities.

### Range expansion of forest-associated taxa drive loss of community variation

Because soil bacterial communities in the forest tended to show high taxonomic overlap with the burned and plantation sites, we asked whether homogenization might rather be driven by changes to the relative abundance of certain taxa. We used DESeq2 – a generalized linear model with a negative binomial distribution-to identify “bloomer” taxa (*i.e*. those whose relative abundance significantly increased by land type). This approach identified 127 taxa that were differentially enriched in the DNA-inferred communities of the burned site relative to the forest (comprising on average 23.85 +/− 9.5% of the DNA-inferred burned site community, and 6.43 +/− 2.6% of the DNA-inferred forest site community), and 192 taxa that were enriched in the plantation relative to the forest (comprising on average 26.89 +/− 10.3% of the DNA-inferred plantation site community, and 5.45 +/− 2.2% of the DNA-inferred forest site community). We removed these bloomer taxa from the community matrices, equalized sampling extent across samples (as described above), and re-calculated pairwise similarity levels within land types. The removal of these taxa from the burned site DNA-inferred community matrix rendered the communities less similar (0.268 +/− 0.005 vs 0.301 +/− 0.006, F_2,196_ = 6.95, *p* = 0.001) and indistinguishable from the forest levels of similarity (0.268 +/− 0.011, Tukey’s HSD *p*_adj_ = 0.999), indicating that their relative abundances are indeed contributing to the increased pairwise similarity of these communities. This was also the case in the plantation, where the removal of the bloomer taxa from the DNA-inferred community matrix also rendered the communities less similar (0.253 +/− 0.005 vs 0.288 +/− 0.006, F_2,184_ = 6.15, *p* = 0.003) and indistinguishable from the forest levels of similarity (0.268 +/− 0.011, Tukey’s HSD *p*_adj_ = 0.300), further supporting the idea that these taxa are driving the increase in levels of pairwise similarity in impacted sites. Beyond decreasing levels of community variation, the bloomer taxa also collectively showed a wider spatial distribution in the sites in which they were more abundant (Burn: freq_burn bloomers in for_ = 0.597 +/− 0.034, freqburn bloomers in burn 0.845 +/− 0.020; Plantation: freq_plantation bloomers in for_ = 0.484 +/− 0.027, freq_plantation bloomers in plantation_ = 0.830 +/− 0.014).

When we test whether these taxa are driving the changes to spatial turnover, however, we do not detect a significant spatial signal for the burned (Mantel r = 0.345, p = 0.08) or plantation (Mantel r = 0.185, p = 0.194) sites, indicating that the weakening or loss of community spatial structure may be driven by additional factors.

The same suite of analyses yielded similar findings for the RNA-inferred communities. Bloomer taxa in the RNA-inferred community comprised on average 35.6 +/− 14.2% of the burned site RNA-inferred community (6.92 +/− 4.0% of the forest site community), and 37.54 +/− 10.6% of the plantation site RNA-inferred community (9.96 +/− 4.1% of the forest site community). As described above, we tested whether the bloomer taxa were contributing to the increased levels of pairwise similarity of the burned and plantation site. Similar to the DNA-inferred findings, the removal of bloomer taxa from the burned site RNA-inferred community matrix rendered the communities less similar (0.306 +/− 0.004 vs 0.327 +/− 0.003, F2,219 = 12.95, *p* < 0.001) and indistinguishable from the forest levels of similarity (0.290 +/− 0.008, Tukey’s HSD *p*_adj_ = 0.124), indicating that their relative abundances contribute to the increased pairwise similarity of these communities. This was also the case in the plantation, where the removal of the bloomer taxa from the RNA-inferred community matrix rendered the communities less similar (0.311+/− 0.004 vs 0.340 +/− 0.004, F_2,219_ = 22.96, *p* < 0.001), but in this case similarity levels were still distinguishable from the forest levels of similarity (0.290 +/− 0.008, Tukey’s HSD *p*_adj_ = 0.029). The RNA-inferred bloomer taxa also collectively showed a wider spatial distribution in the sites in which they were more abundant (Burn: freq_burn bloomers in for_ = 0.566 +/− 0.04, freq_burn bloomers in burn_ = 0.860 +/− 0.022; Plantation: freq_plantation bloomers in for_ = 0.506 +/− 0.03, freq_plantation bloomers in plantation_ = 0.854 +/− 0.015), but when we test whether these taxa are driving the changes to spatial turnover we do not detect a significant spatial signal for the burned (Mantel r = 0.006, p = 0.474) or plantation (Mantel r = 0.413, p = 0.082) sites following their removal. Thus, the identification of bloomer taxa in the DNA- and RNA-inferred communities has helped to identify the fraction of the community that is contributing to higher levels of community pairwise similarity.

## DISCUSSION

Conversion of tropical rainforest to agriculture is one of the leading drivers of biodiversity loss and biotic homogenization worldwide (1–4). Gaining a better understanding of the mechanisms driving biotic homogenization is a priority if we are to predict or mitigate changes to communities or their ecosystem functions (8, 9). We used a spatially explicit design across a chronosequence of land use change in the Congo Basin to investigate mechanisms of community homogenization. We used two windows into the structure of soil prokaryotic communities: 1) 16S rRNA (RNA) community inference – which should enrich for the active fraction of the community, and 2) 16S rRNA gene (DNA) community inference – which includes both active and inactive members, as well as “relic” DNA from dead cells (82, 83). Our results fit into a broader context of other studies that emphasize the importance of using RNA alongside DNA to investigate the impacts of environmental change on microbial communities (34, 35).

Ecosystems can develop spatially autocorrelated environmental conditions (*i.e*. a distance-decay in environmental similarity) through a combination of localized physical forces or community processes (84). Slash-and-burn conversion in our system appears to disrupt this spatial structure, while introducing variation. This form of conversion is a relatively uniform type of disturbance, in that all the aboveground vegetation gets removed and burned across the landscape, which likely drives the loss of spatial structure of the soil environment. The intensity of fire across a landscape, however, is often patchy, depending on certain local factors such as, *e.g*., the amount of biomass, or levels of moisture. Thus this form of disturbance could introduce environmental variation that shows little coherent spatial structure. This insight is important when we consider the relationship between community structure and the environment.

Communities can be homogenized by two main mechanisms: 1) the homogenization of the environment driving convergence of communities (12, 13), or 2) increased biotic mixing, driven by the breakdown of dispersal barriers and/or the range expansion of previously present taxa (6, 10, 14, 15, 85, 86). If community homogenization is driven by environmental homogenization, community turnover should continue to track environmental turnover, even when spatial structure is lost. We see this in our data when we infer community structure using RNA, but not DNA, suggesting that environmental spatial homogenization is likely a strong driver of the spatial homogenization of the RNA-inferred community. The decoupling of responses in the RNA- and DNA-inferred communities could represent differing levels of contribution from homogenization mechanisms. Our results suggest that taxa that are enriched in the burned or plantation sites relative to the forest are contributing to the loss of community variation (i.e. average pairwise dissimilarity) in those sites. Those taxa also collectively show wider spatial distributions (i.e. higher occurrence frequencies) in the disturbed sites relative to the forest. These findings are consistent with the idea of a range expansion, and the fact that we saw this trend in both the RNA- and DNA-inferred communities suggests that identifying this type of homogenization mechanism may not require RNA-based community inference. A similar pattern has been observed in Amazonian sites that have undergone conversion to cattle pasture, where prokaryotic taxa shared across forest and agricultural sites tended to be more widespread in the agricultural sites (6), and fungal communities in agricultural sites tended to be enriched in generalist taxa that were more widespread (15). Thus by distinguishing communities using RNA and DNA, we see that only part of the community seems to be responding to the environmental changes associated with conversion, while communities inferred via both methods appear be shaped by biotic factors such as the breakdown of dispersal barriers and/or the range expansion of certain taxa.

The use of 16S rRNA as a proxy for activity has been the subject of recent controversy. Of particular concern are two main issues: the assignment of false positives (i.e. dormant taxa misidentified as active (28)), and the inaccurate assessment of activity levels (e.g. driven by comparing ratios of the relative abundance of taxa in the RNA- vs DNA-inferred communities (29–32)). The ribosomal content of a community, however, should be at least enriched with the taxa that are active and/or growing, and there are a number of studies that support the notion that rRNA-inference represents activity. For example, if the active fraction of a community is more likely to be interacting with the environment than the dormant fraction (which is likely avoiding the current environmental conditions), then we would expect a stronger correspondence between environmental conditions and community turnover in a community that is enriched in active taxa (19). Indeed this has been shown both along a marine environmental gradient (33) and a grassland soil system experiencing re-wetting following drought (34). It has also been shown that N-addition to forest soil elicits a stronger response in communities inferred from 16S rRNA than rDNA (35). Our results contribute to this narrative by showing that RNA-inferred community turnover persistently tracks environmental turnover, while this association is lost when inferring only with DNA. We also see that the RNA-inferred community shows a more pronounced loss of community variation and spatial structure than the DNA-inferred community. Thus while rRNA inference may have certain limitations, our results, alongside others, suggest that this method should be enriching for active taxa, and this can have important implications for both qualitative and quantitative conclusions, especially in systems with strong environmental gradients.

Tropical ecosystems are characterized by immense heterogeneity, and this could make the task of detecting general responses to land use change difficult. Two important steps towards gaining a better understanding of common microbial responses to tropical land use change include 1) expanding the breadth (i.e. the geographic representation) of regions sampled, and 2) increasing the resolution of our study systems (e.g. by including more sites along the conversion continuum). Our study allows us to ask whether commonalities exist between our findings and those reported from other tropical ecosystems undergoing land use change. The changes we see to the spatial structuring of communities (i.e. a diminished distance-decay relationship) are consistent with responses reported from the Amazon Basin (6, 25). While our study was not replicated at the land type level-restricting our level of inference regarding how representative our findings are of other Congo Basin areas-our results at least suggest that a diminished rate of community distance-decay may be common across tropical areas facing a similar threat. The method of conversion may be driving this similarity in microbial community response. The predominant method for converting tropical rainforests to agriculture is the use of slash-and-burn techniques (87). By including a recently slash-and-burned site in our design, we have gained a rare glimpse into the impacts directly following the initial step in agricultural conversion. Already at this stage we see that the loss of community spatial structure (*i.e*. distance-decay) has occurred. What this suggests is that, at least initially, spatial homogenization can be driven by the act of conversion, rather than other management practices such as planting or crop choice. Thus by targeting a region that has otherwise not been sampled, and increasing the resolution by which we survey the conversion process, we have gained new insights that may help to elucidate common community responses to tropical land use change.

Considering the rate and magnitude by which tropical rainforests are being converted to agriculture (4), gaining a mechanistic understanding of community responses to environmental change is imperative (9). Future efforts could investigate whether the functional potential (i.e. gene content) or trait distributions of a community are similarly impacted by land use change (37, 88), or whether ecosystem functions (*e.g*. those involved in nutrient cycling or greenhouse gas emissions) are impacted by community homogenization. Our work highlights the importance of distinguishing between metabolic states of microbial community members, if we are to better understand community responses to environmental change. Lastly, our work demonstrates that trends in our system are consistent with those reported from geographically disparate areas (e.g. the Amazon Basin), suggesting that despite large differences between these areas, land use change may drive predictable community changes.

## ACKNOWLEDGEMENTS

We thank the Government of Gabon, Centre National de la Recherche Scientifique et Technologique for permission (Permit N^o^ 866/MENESTFPRSCJS/CENAREST/CG/CAB) to conduct this study. We also thank P. Voua Otomo and A. Litona Boubeya for their help in collecting samples. The Smithsonian Conservation Biology Institute, the Gabon-Oregon Transnational Center on Environment and Development, and Shell Gabon provided financial and logistical support.

